# Coronavirus Nucleocapsid Proteins Hijack Host Protein Kinase A Catalytic Subunit α into Nucleus to Evade from STAT1 Signaling

**DOI:** 10.64898/2026.01.03.697510

**Authors:** Jidong Xu, Yuting Xiao, Wanqing Lu, Ziyao Song, Lei Zhou, Qin Gao, Xiaohan Xu, Ying Shan, Weihuan Fang, Lingyan Zhao, Xiaoliang Li

## Abstract

The role of the cAMP/PKA pathway in antiviral innate immunity, including during coronavirus infections, remains unclear. We discovered coronavirus N proteins initiate cAMP-ADCY10-PKA cascade. Cytoplasmic PKA activates STAT1 independently of canonical JAK/TYR2 signaling. Coronavirus N proteins, via a conserved arginine (e.g., PEDV R58, SARS-CoV-2 R92), directly bind and sequester PKA Cα into the nucleus to evade STAT1 activation. Using PEDV as a model, mutant viruses with N^mutNLS^ and N^R58A^ were generated. Wildtype PEDV infection suppressed STAT1 activation in cells expressing PKA Cα/PKA Cα^S339A^ deficient in STAT1 phosphorylation. However, in rXS0101^mutNLS^- or rXS0101^R58A^-infected cells, only PKA Cα^S339A^ expression inhibited STAT1 activation. Downregulation of STAT1 activation is accompanied with increased viral replication. This study first elaborates that PKA Cα activates STAT1 in the cytoplasm of infected cells distinctly from canonical JAK/STAT1 signaling, while coronaviruses evade this antiviral response by sequestering PKA Cα into the nucleus via direct N protein interaction.

## Introduction

Coronaviruses (CoVs) belong to the family *Coronaviridae* within the order Nidovirales and can be divided into the four genera α-, β-, Δ-, and γ-coronavirus. These CoVs differ in their cellular/tissue tropism and their ability to infect different mammalian species including humans. So far, more than fifteen coronavirus-related diseases have been described in animals and seven, in humans^1^. Severe acute respiratory syndrome coronavirus 2 (SARS-CoV-2), the causative agent of the coronavirus disease 2019, is an emerging zoonotic β-CoV that has spread all over the world causing high mortality and critical social issues in the past several years^2,3^. Porcine epidemic diarrhea virus (PEDV), belonging to α-CoV, is a highly contagious pathogen in pigs, leading to watery diarrhea, vomiting, dehydration and even death in suckling piglets^4,5^. The genomes of all coronaviruses are arranged similarly, and encode 16 nonstructural proteins (Nsp 1-16) and 4 structural prh h hoteins (spike protein, S; envelop protein, E; membrane protein, M; and nucleocapsid protein, N) as well as an accessory protein (ORF3) in PEDV^6,7^ or several others in SARS-CoV2^8,9^.

The N protein is well conserved in all coronaviruses and multifunctional with effects on virus life cycle (RNA synthesis, RNA packing, viral assembly, etc.) as well as host cell cycle regulation, cell stress response, evasion of innate immunity and signal transduction^10–14^. It is, structurally, composed of 3 conserved functional domains: N-terminal Domain (NTD) for RNA binding, C-terminal Domain (CTD) for RNA binding and dimerization, and the linker region between NTD and CTD^15,16^. SARS-CoV-2 N protein promotes NLRP3 inflammasome activation by direct interaction with NLRP3 and induces severer lung injury^17^. SARS-CoV2 also employs its N protein to antagonize type I interferon (IFN-I) expression by interacting with TRIM25 and inhibiting RIG-I signaling^18^, and to block type I IFN signaling by suppressing STAT1/2 phosphorylation and nuclear localization^19^. PEDV N protein is also confirmed as a key modulator of host immune responses, such as inhibition of nuclear translocation of NF-κB to suppress type III IFN production^20^ and induction of the S-phase arrest of host cells to favor viral replication^11^. Subcellular localization of CoVs’ N, including those of SARS-CoV, SARS-CoV2 and PEDV, is determined by its nuclear localization sequence (NLS)^21,22^. Shi et al. reported that PEDV N protein contains NLS and NES (nuclear exporting signal) that mediate its translocation into/out of the nucleus during PEDV infection^23^. Xu et al. further identified an NLS (^261^-PKKNKSR-^267^) for PEDV N entry into the nucleus where it repressed HDAC1 (histone deacetylase 1) transcription and downstream STAT1 activation^24,25^. The above studies indicate that the coronavirus N proteins serve multiple functions in the infected cells. Elucidating their underlying mechanisms will help better understanding of the pathogenesis of these highly contagious and pathogenic coronaviruses.

The 3′-5′-cyclic adenosine monophosphate (cAMP), an important intracellular second messenger, plays versatile roles in signaling transduction and cellular responses^26,27^. Transducing signals from G-protein coupled receptors (GPCRs) at the cell membrane lead to significant changes of the cAMP level that affects an array of cellular responses involved in cell proliferation and differentiation, glucose and lipid metabolism, gene expression, immune responses, and others^28–30^. In mammalian cells, cAMP is transformed from cellular ATP by the adenylate cyclase (ADCY) family members. There are 9 transmembrane ADCYs (ADCYs1-9) and 1 soluble ADCY (ADCY10) identified so far^31^. ADCY10 is conserved among different species and distributed in cytoplasm, nucleus and mitochondria^32,33^. Different from ADCYs1-9 which are activated by GPCRs, ADCY10 could be regulated by the cellular proteins^34^, leading to changes of the cAMP level in the cytoplasm and activation of a series of protein kinases, including protein kinase A (PKA)^26^. PKA holoenzyme is a hetero-tetrameric protein comprised of two regulatory subunits and two catalytic subunits that targets serine and threonine residues^35^. Upon binding to cAMP, the regulatory subunits undergo conformational changes and dissociate from the catalytic subunits available for phosphorylating the downstream effectors^36^. While the cAMP/PKA pathway is a crucial mediator in quite a number of biological functions, including several virus infections such as hepatitis C virus, human immunodeficiency virus and influenza A virus^37–39^, whether this pathway is involved in coronavirus infection is unknown.

Our initial experiments revealed that PEDV infection resulted in marked increase of the cellular cAMP level. We then wondered which ADCY is activated, which viral protein is involved and what would be the downstream effects and mechanism(s) thereof on the virus. We found that ADCY10 is the enzyme responsible for PEDV-induced cAMP upregulation and PKA Cα phosphorylation. We further identified that PKA Cα phosphorylates STAT1, distinct from JAK, to activate IFN-λ3 signaling. However, coronaviruses utilize their N proteins to hijack PKA Cα into the nucleus via direct interaction. This led to diminished abundance of PKA Cα in the cytosol and downregulation of PKA Cα-STAT1-mediated antiviral innate immunity.

These novel findings reveal that PEDV, even other coronaviruses, might evade from the host antiviral IFN responses by deploying their N proteins to dampen the PKA signaling and STAT1 activation.

## Materials and methods

### 1. Reagents and antibodies

The chemical regents and antibodies used in this study were as follows: sp-8-CPT-cAMP (8-Bromo, 116818, MERK), H 89 2HCl (S1582, Selleck), HL-362 (also termed Forskolin, S2449, Selleck), SQ22536 (S8283, Selleck), Ruxolitinib (S1378, Selleck); PKA Cα antibody (#4782, CST), Phospho-PKA C (Thr197) (D45D3) rabbit mAb (#5661, CST), STAT1 (D1K9Y) rabbit mAb (#14994, CST), Phospho-STAT1-Y701 rabbit mAb (AP0054, ABclonal, China), ISG15 rabbit pAb (A1182, ABclonal), OAS1 rabbit pAb (A2530, ABclonal), β-actin mouse mAb (MA5-15739, ThermoFisher), GAPDH mouse mAb (AF0006, Beyotime, China), histone H3 mouse mAb (AF0009, Beyotime), HA-tag rabbit mAb (#3724, CST), Flag-Tag rabbit mAb (#14793, CST), PEDV N mouse mAb (produced and reserved in our laboratory); goat anti-mouse IgG (Invitrogen) and goat anti-rabbit IgG (Invitrogen); Alexa Fluor 488 goat anti-rabbit IgG (Invitrogen), and Alexa Fluor 555 donkey anti-mouse IgG (Invitrogen).

### 2. Cell lines and virus stock

The porcine intestinal epithelial cell line (IPEC-J2) and VERO-E6 cell line were maintained in our laboratory. IPEC-J2 cells were cultured in Dulbecco’s Modified Eagle Medium/Nutrient Mixture F-12 (DMEM/F12, Gibco) supplemented with 10% fetal bovine serum (FBS, Gibco), 100 U/mL of penicillin, 0.1 mg/mL of streptomycin, and 0.25 mg/mL of amphotericin B (Gibco) at 37°C with 5% CO_2_. The VERO-E6 cells were cultured in DMEM medium supplemented with 10% FBS, 100 U/mL of penicillin, 0.1 mg/mL of streptomycin, and 0.25 mg/mL of amphotericin B at 37°C with 5% CO_2._

The PEDV strain ZJ15XS0101 (GenBank accession No. KX550281.1) used in this study was previously isolated from a clinically diseased pig in Zhejiang Province, China^40^. The virus was propagated for 27 passages and titrated in the VERO-E6 cells before use for viral infection. For the recombinant PEDV rXS0101, rXS0101-mutNLS, and rXS0101-R58A construction, an infectious cDNA plasmid pBAC-XS0101 was constructed using the wildtype XS0101 strain as the parent and the specific primer pairs (Table 1). The positive clones were used to amplify the full-genome sequence of PEDV in 6 segments, and the PCR products were subjected to agarose gel electrophoresis for verification. The pBAC-XS0101 plasmid or those with mutated NLS (mut-NLS) or R58A substitution were then transfected into 293T cells, and the cell cultures were harvested at 36 h post transfection. The cultures underwent freeze-thaw cycles and centrifugation, and the supernatant samples were then used to infect Vero cells to generate rXS0101, rXS0101-mutNLS and rXS0101-R58A which were propagated upon verification by sequencing, titrated and utilized for viral infection assays.

**Table. 1.**
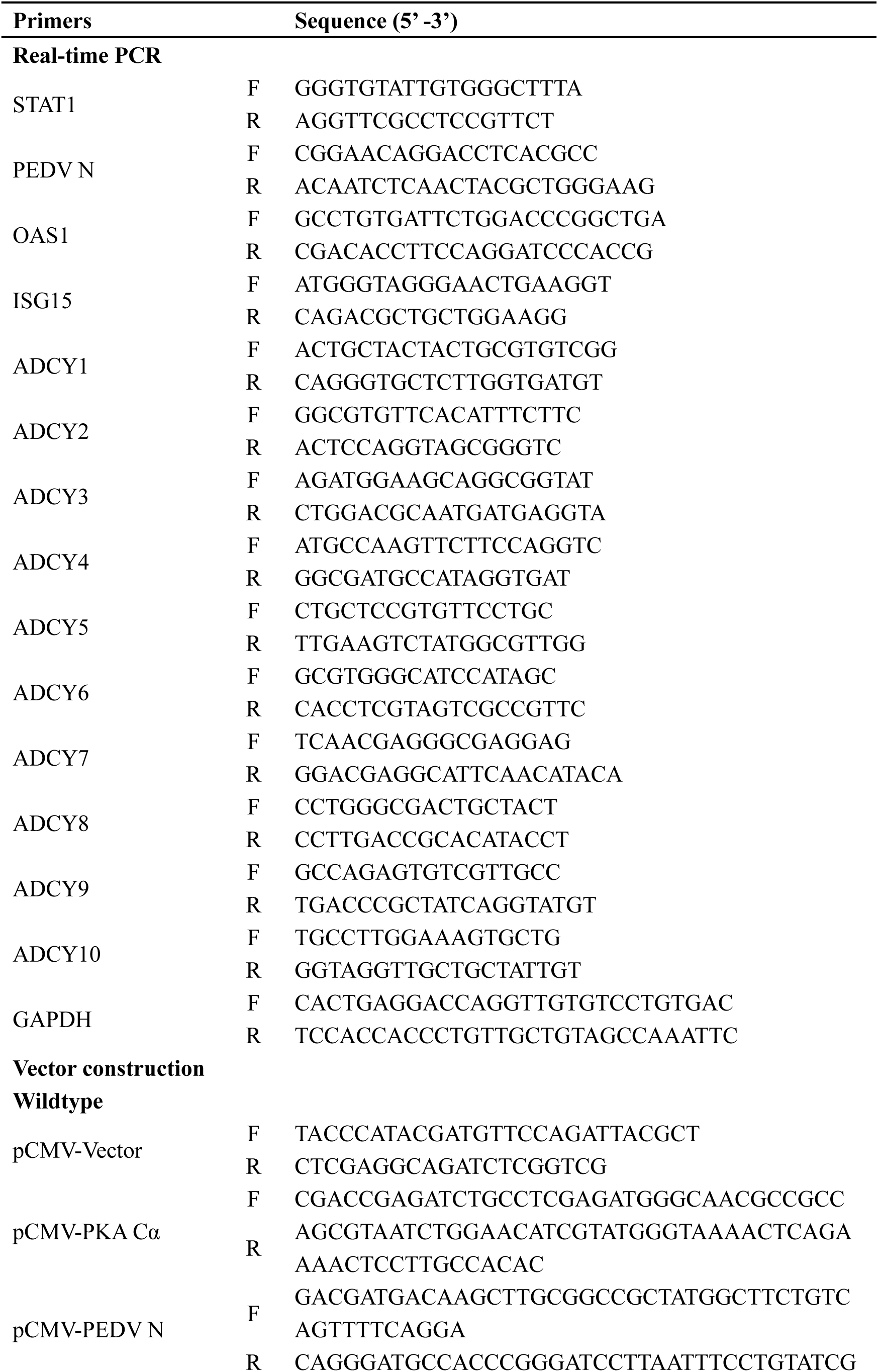
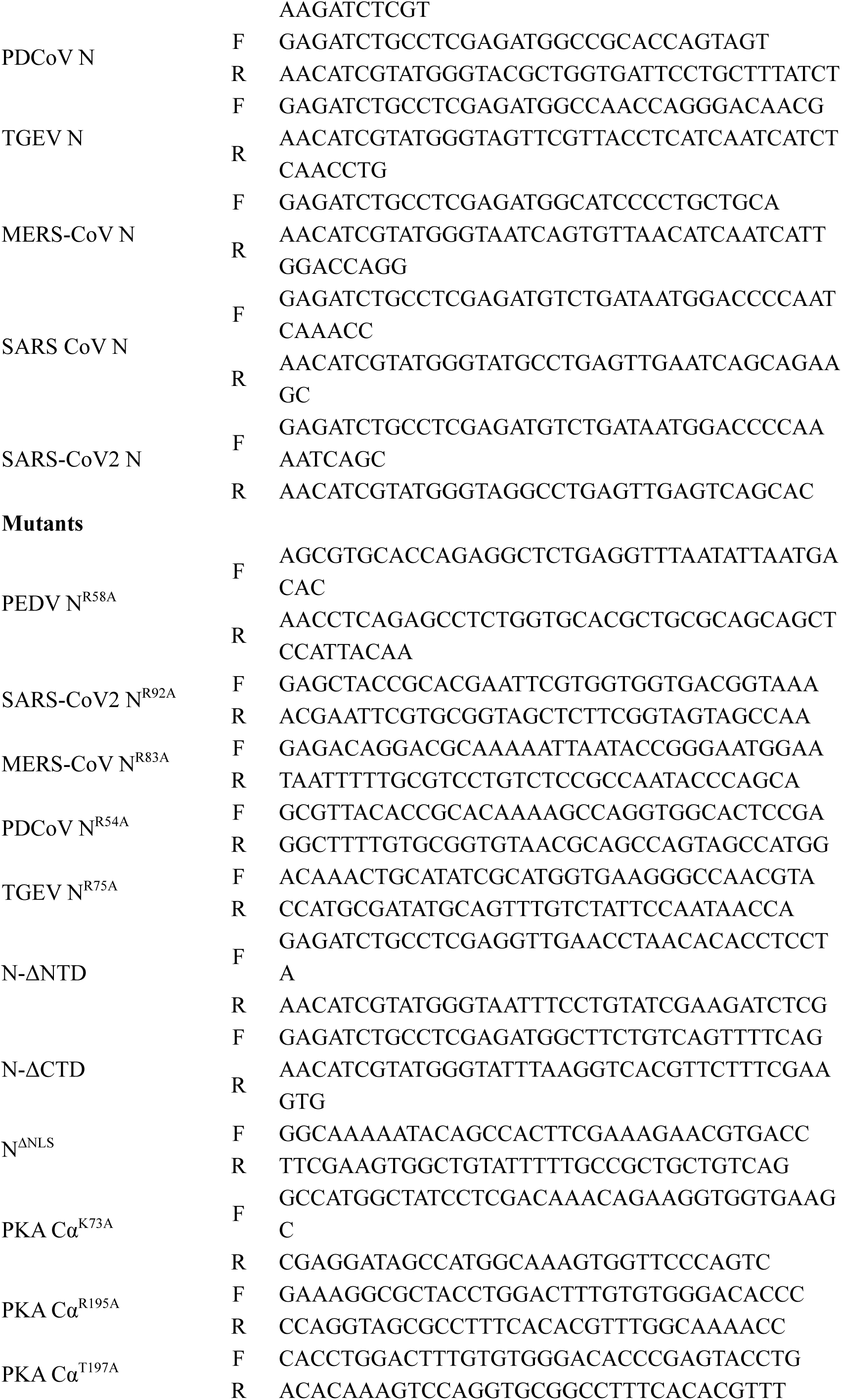
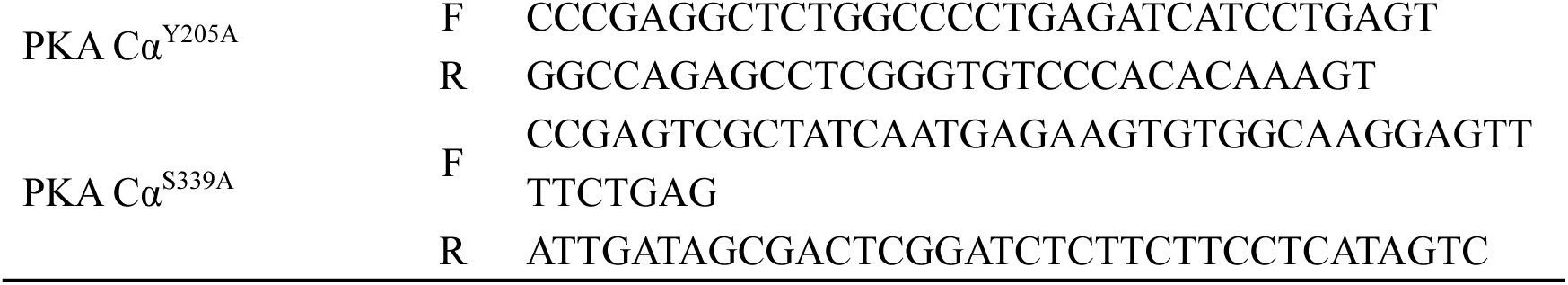
Primers used in this study.

### 3. Recombinant plasmids and transfection

Full-length porcine PKA Cα sequence (GenBank accession No. XM_003123353.4) was cloned from the cDNA template of IPEC-J2 cells using specific primer pairs (Table 1) and inserted into the pCMV-based plasmid with an HA tag. The recombinant plasmids expressing coronavirus N proteins were constructed with HA tag/Flag tag using the same method as above with specific primer pairs shown in Table 1. Different N protein mutants (PEDV N^R58A^, SARS-CoV2 N^R92A^, MERS-CoV N^R83A^, PDCoV N^R54A^, TGEV N^R75A^, N-ΔNTD, N-ΔCTD, N-ΔNLS) and PKA Cα mutant (PKA Cα^T197A^) were constructed using site-directed mutagenesis or truncations, respectively, with specific primers (Table 1).

For protein overexpression or gene knockdown with small interfering RNA, the recombinant plasmids or siRNAs were transfected into IPEC-J2/Vero-E6 cells using Lipofectamine 3000 (Invitrogen) according to the manufacturer’s instructions. After transfection for appropriate times, the cells were lysed for RNA isolation or total protein extraction.

### 4. PEDV infection and recombinant IFN-λ3 treatment

For PEDV infection, IPEC-J2 cells were grown to about 60% confluency, after which the cells were infected with the virus at the multiplicity of infection (MOI) of 1.0. After 4 h-adsorption at 37°C and 5% CO2, the inocula were washed off with sterile phosphate-buffered saline (PBS) and replaced with the maintenance medium (DMEM with 4 μg/mL of trypsin). RNA extraction and total protein lysis were performed at the proper time points.

After transfection or PEDV infection, the IPEC-J2 cells were treated with the recombinant IFN-λ3 protein (rIFN-λ3, RP00219, ABclonal) for 24 h at final concentration of 50 ng/mL to activate JAK-STAT1 signaling. Afterwards, the cells were harvested for total RNA and protein extraction.

### 5. Quantitative RT-PCR

Total RNA samples from the above experiments were extracted using the TRIzol^TM^ Reagent (Thermo). The cDNAs were synthesized using the HiScript II Q RT supermix for qPCR (+DNA wiper) (Vazyme, China). Relative transcription levels of *PEDV N*, *ISG15*, *OAS1* and *ADCY1-10* were analyzed by quantitative PCR (qPCR) with specific primer pairs (Table 1) and AceQ universal SYBR qPCR master mix (Vazyme). The qPCR profile was as follows: 95°C for 10 min, 40 cycles of 95°C for 15 s, 58°C for 50 s, and 72°C for 2 s, and the melting curve obtained from 65°C to 95°C. Relative transcription levels of the target genes were normalized to glyceraldehyde-3-phosphate dehydrogenase (*GAPDH*), and analyzed by the threshold cycle (2^-ΔΔCt^) method as described previously^41^.

### 6. Separation of nuclear and cytoplasmic fractions

After PEDV infection or overexpression, the cells were treated with NE-PER nuclear and cytoplasmic extraction reagents (Thermo) to collect cytoplasmic and nuclear proteins. Protein concentrations were determined using the Bicinchoninic Acid (BCA) Protein Assay Kit (Thermo). The target protein levels of PEDV-N and PKA were visualized by Western blotting and quantified by densitometric analysis. GAPDH and Histone H3 were used as the cytoplasmic and nuclear markers, respectively.

### 7. Immunoprecipitation (IP) and Western blotting

The IPEC-J2 cells or VERO-E6 cells were infected with PEDV or transfected with the indicated plasmids for proper times and then harvested for total protein lysis. Except for the IP samples, the cell lysates were first quantified by the BCA protein assay kit. The IP cell lysate samples were incubated at 4°C overnight with anti-HA Affinity Gel Sepharose or anti-Flag Affinity Gel sepharose. The beads containing bound proteins were centrifuged, washed with D-PBS, mixed with the protein loading buffer and denatured on the boiling water-bath.

For Western blotting, the protein samples were loaded onto the 12% SDS-PAGE gel for protein separation. After blotting onto the polyvinylidene difluoride (PVDF) membranes (Millipore), the proteins were probed by the specific primary antibodies described above. The protein bands were visualized with an ECL kit (Cyanagen, Italy) and an imaging system (SageCreation, Beijing, China). Target protein levels were quantified by the ImageJ software (National Institutes of Health, USA).

### 8. Cell staining and confocal microscopy

At 24 h after PEDV infection or transfection, the cells were fixed with 4% paraformaldehyde (Beyotime) at room temperature for 30 min, and treated with 0.1% Triton X-100 (Sangon Biotech, China) to permeabilize the cell membrane. The cell samples were then incubated with primary and secondary antibodies for 1 h. The cell nuclei were labeled with 2-(4-amidinophenyl)-6-indolecarbamidine dihydrochloride (DAPI). At the end of each incubation, the cells were washed thoroughly with D-PBS. Subcellular localization of PKA Cα and PEDV-N proteins was visualized under the laser scanning confocal microscope (Olympus #IX81-FV1000).

### 9. cAMP detection assay

To measure the cytoplasmic cAMP levels, the IPEC-J2 cells or the VERO-E6 cells were infected with PEDV at different MOIs, or transfected with the plasmids carrying different CoV N proteins for appropriate time periods. The cells were harvested and lysed for cAMP measurement with a cAMP ELISA Detection Kit (Genscript, China) according to the manufacturer’s instructions.

### 10. Statistical analysis

Each experiment in this study was repeated at least 3 times. Relative mRNA transcripts by qPCR or relative protein levels from densitometric analysis were analyzed in GraphPad Prism 9 software, and shown as means ± standard deviations (SD). Significant differences were determined by the two-tailed student’ *t* tests (\**P*<0.05, \*\**P*<0.01 and \*\*\**P*<0.001).

## Results

### 1. Coronaviruses employ N proteins to upregulate ADCY10, leading to increased cytoplasmic cAMP and PKA Cα activation

Our initial experiments indicated that PEDV infection increased cellular cAMP in an MOI-dependent manner (**Fig. 1A**). cAMP is known to regulate the transcription of a variety of target genes for different biological processes by activating PKA^28^. Therefore, we examined the phosphorylation status of PKA Cα, the active form of PKA. PEDV infection did promote PKA Cα phosphorylation in IPEC-J2 cells similar to treatment with the cAMP analogue sp-8-CPT-cAMPS (**Fig. 1B/b**). Subsequently, we attempted to examine which PEDV protein(s) are involved in cellular cAMP production. Of the PEDV proteins tested, only the N protein could significantly induce cAMP production in IPEC-J2 cells (**Fig. 1C**). Similar to PEDV infection, the N protein overexpression could also induce PKA Cα phosphorylation in IPEC-J2 cells (**Fig. 1D/d**). These results suggest that PEDV might activate the cAMP-PKA signaling by employing its N protein.

**Fig. 1.**
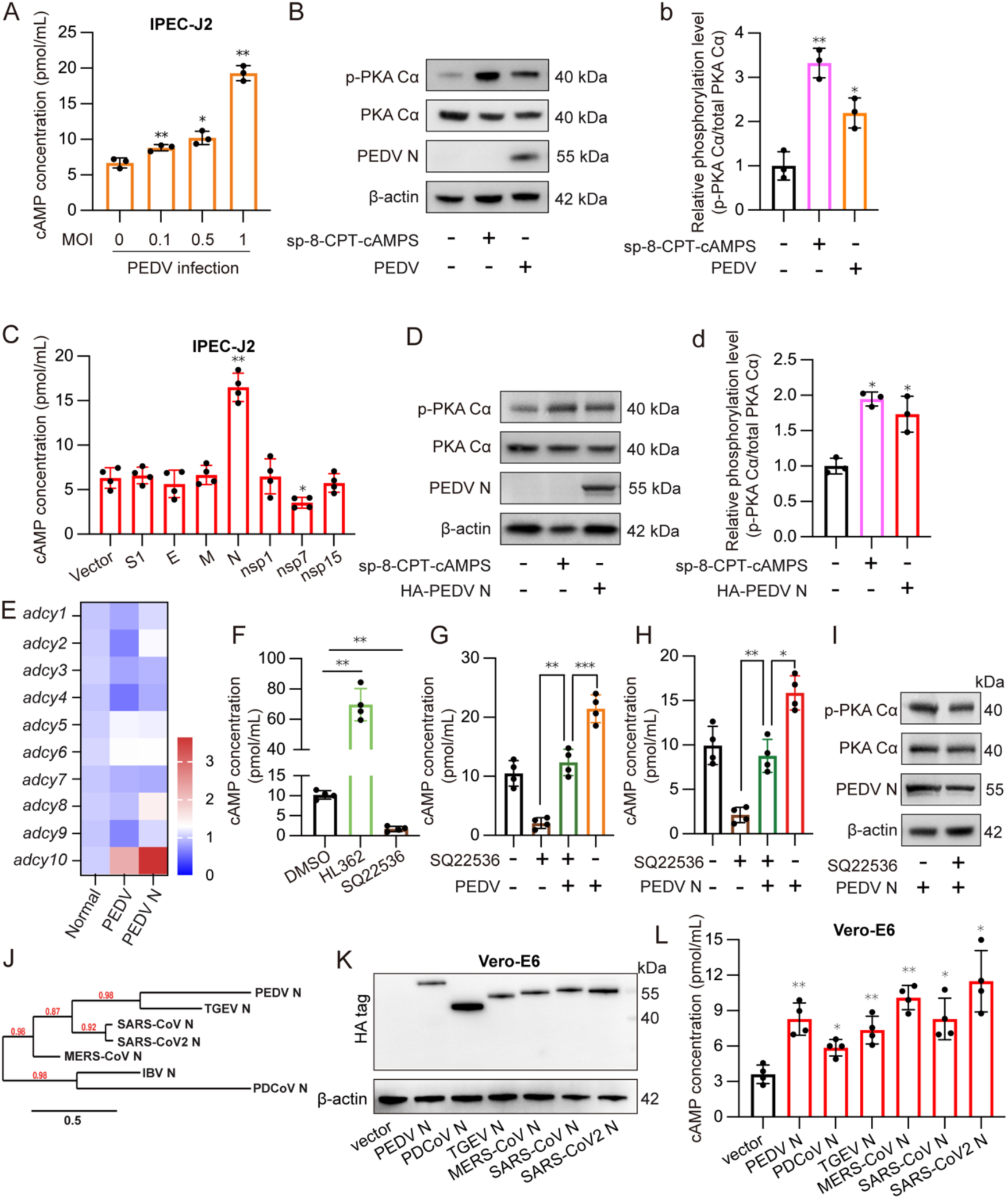
Coronaviruses use the nucleocapsid proteins to enhance cAMP production and PKA activation by upregulating soluble adenylyl cyclase 10. (A) IPEC-J2 cells were infected with PEDV at the MOI of 0.1, 0.5, and 1. At 24 hpi, the cell lysates were extracted for cAMP detection by an ELISA kit. (B) Detection of total PKA Cα, p-PKA Cα and PEDV N levels in the cells infected with PEDV (MOI 1 for 24 h) or treated with 10 μM of cAMP analogue sp-8-CPT-cAMPS for 30 min (positive control). (b) Density analysis of (B) from 3 independent repeats. (C) cAMP levels in IPEC-J2 cells expressing PEDV proteins (S1, E, M, N, nsp1, nsp7, and nsp15, in pCMV-HA tagged plasmids). (D)&(d) Same as (B)&(b) but with viral N protein expression for 24 h. (E) Heat map showing the transcription levels of different *ADCY* genes after PEDV infection at the MOI of 1 for 24 h, or PEDV N expression for 24 h in IPEC-J2 cells. (F) cAMP levels in the IPEC-J2 cells treated with HL362 or SQ22536 for 24 h. (G-H) The cells were pretreated with SQ22536 for 30 min, and then infected with PEDV for 24 h (G), or transfected with pCMV-PEDV N for 24 h (H). (I) Detection of total PKA Cα and p-PKA Cα in the N expressing cells of (H). (J) Phylogenetic tree showing genetic relationship of different coronavirus N proteins. (K) Vero-E6 cells were transfected with pCMV-HA tagged plasmids containing N sequences of different coronaviruses for 24 h and then lysed Western blotting with anti-HA antibody. (L) cAMP levels in the Vero-E6 cells expressing different coronaviruses’ N proteins. The results of panels A, b, C, d, F, G, H, and L were obtained from at least 3 independent experiments, and were shown as means ± SD: *, *p*<0.05; **, *p*<0.01; ***, *p*<0.001.

Cellular cAMP production is predominately regulated by adenylate cyclases (ADCYs)^42^, and there are ten ADCYs (ADCY1-10) identified in pigs. In order to investigate which ADCY is responsible for cAMP generation, transcriptional analysis of the *adcy* genes was conducted in PEDV-infected or N-protein expressing IPEC-J2 cells. **Figure 1E** shows that *adcy10*, encoding the only soluble ADCY protein, rather than the other *adcy* genes (*adcy1*-*adcy9*), was upregulated by both PEDV infection and PEDV N overexpression. To further confirm that cAMP induction by PEDV N protein is ADCY10-dependent, we employed the specific ADCY inhibitor, SQ22536 (**Fig. 1F** vs HL362 as activator), to see if increased cAMP in PEDV infected or PEDV N expressing cells could be repressed. We found that the PEDV- and its N protein-induced cAMP production could be markedly restrained by SQ22536 (**Fig. 1G & 1H**). Similarly, N protein-induced PKA Cα phosphorylation was inhibitable by SQ22536 (**Fig. 1I**). More importantly, all N proteins from representative coronaviruses, including SARS-CoV2, could induce cAMP production in the Vero-E6 cells (**Fig. 1J-1L**) and upregulate transcriptional levels of the *adcy10* gene only (**Fig. S1**). These results clearly indicate that the coronaviruses employ their nucleocapsid proteins to induce ADCY10-mediated cAMP production and PKA activation.

### 2. PEDV N protein promotes PKA Cα nuclear localization in IPEC-J2 cells

Studies from our lab and others indicate that CoV N proteins could shuttle between cytoplasmic and nuclear compartments^21,22,24,43^, while the PKA catalytic subunits could be localized to the nuclear compartment by interaction with A-kinase-interacting protein (AKIP1) in forskolin-stimulated HeLa cells^44^, and with A-kinase anchoring proteins 6 and 8 (AKAP6 and AKAP8)^30^. We speculated if PEDV N could be involved in nuclear translocation of PKA Cα. While PKA Cα was distributed in both cytoplasm and nucleus under normal conditions, PEDV N expression significantly promoted nuclear distribution of phosphorylated and non-phosphorylated forms of PKA Cα in addition to increase of its phosphorylation in the total cell lysates (**Fig. 2A/a & 2B/b**). Confocal imaging also revealed its preponderance in the nuclear compartment in PEDV-infected cells (enriched in the multi-nuclear syncytia), as compared with the mock, (**Fig. 2C**). In PEDV-infected cells overexpressing PKA Cα, both the N protein and PKA Cα were enriched in the nuclear compartment, as compared with either treatment alone (**Fig. 2D**), suggesting their interdependence towards nuclear localization.

**Fig. 2.**
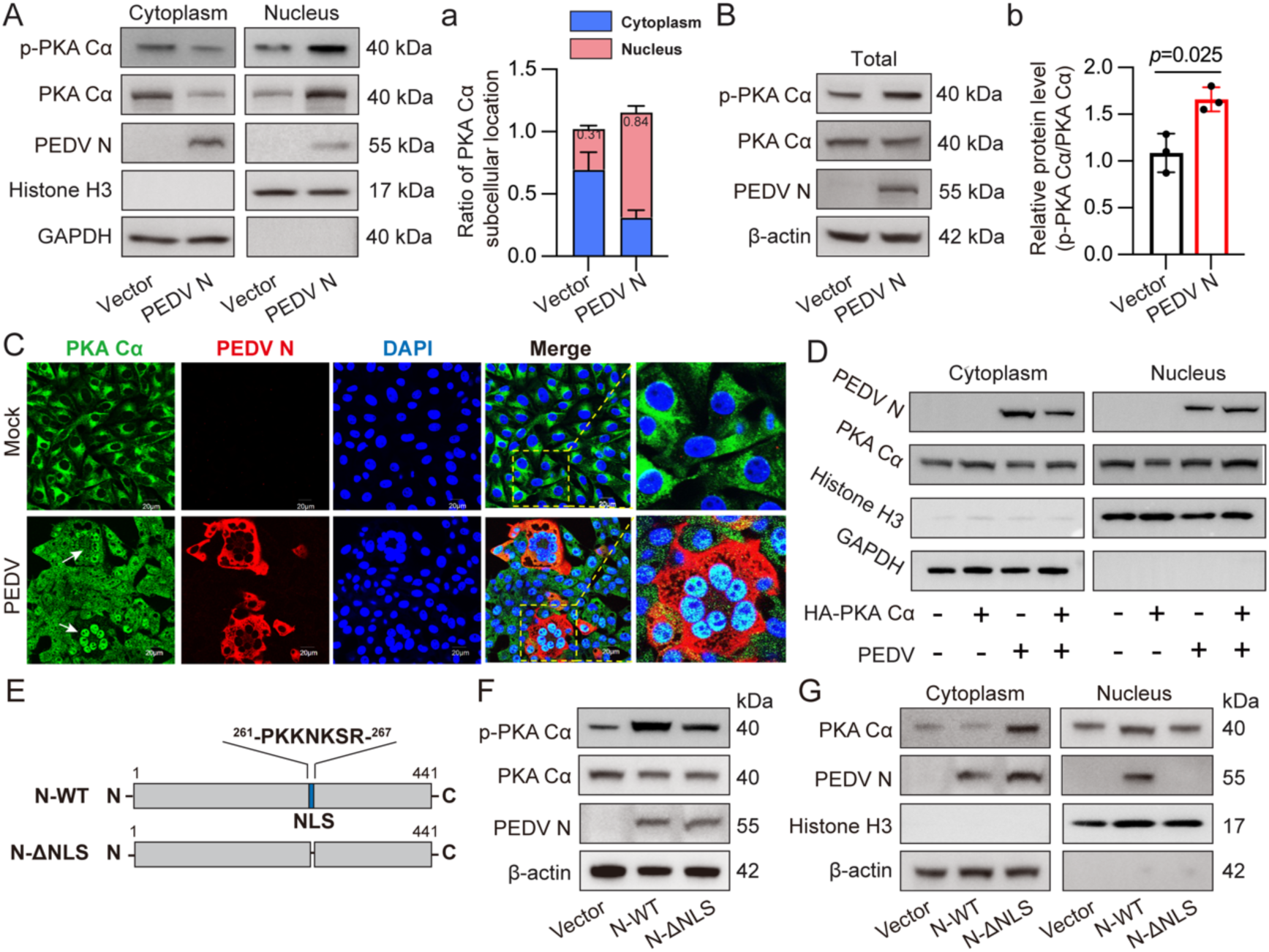
Nuclear localization signal sequence is required for the PEDV nucleocapsid protein to enhance PKA Cα entry into nuclei. (A) & (B) Detection of total and phosphorylated PKA Cα in the nuclear and cytoplasmic fractions and total cell lysates of the IPEC-J2 cells expressing PEDV-N. (a) & (b) Densitometric analysis of (A) and (B). (C) Immunochemical visualization of PKA Cα in nuclei of the multinuclear syncytia of PEDV-infected IPEC-J2 cells. (D) Subcellular distribution of PEDV N and total PKA Cα in the IPEC-J2 cells with PKA Cα overexpression for 24 h followed by PEDV infection. (E) Schematic diagram showing construction of PEDV N mutant with deletion of NLS. (F) & (G) PKA Cα phosphorylation and its subcellular distribution in the IPEC-J2 cells expressing wildtype PEDV N protein and its NLS deletion mutant. The experiments of panels A and B were repeated 3 times for density analysis (a and b). The results of panels a and b were shown as means ± SD.

Our previous study found that the PEDV N protein contains a putative NLS (^261^-PKKNKSR-^267^) within the NTD region that is involved in its nuclear localization and the NLS deletion mutant N-ΔNLS is present only in the cytoplasm (**Fig. 2E**)^24^. Here, we observed that NLS deletion still enhanced PKA Cα phosphorylation similar to its wildtype N (**Fig. 2F**), but led to downregulation of PKA Cα in the nuclear fraction (**Fig. 2G**), indicating that nuclear translocation of PEDV N protein is required to promote PKA Cα entry into the nucleus. This also suggests that there is interaction between the two proteins.

### 3. The conserved arginine residue in the NTD of coronavirus N proteins is responsible for binding to and nuclear translocation of PKA Cα

The preceding findings indicate that PEDV utilizes its N protein to regulate PKA Cα nuclear localization. We then explored if CoV N proteins interact with PKA Cα. Co-IP either with anti-HA-PKA Cα or with anti-Flag-PEDV N revealed binding of the two proteins (**Fig. 3A & 3B**). To determine the region of the PEDV N protein in binding to PKA Cα, the truncated N mutants N-ΔCTD and N-ΔNTD were made based on the generic structural features of the coronaviruses, including PEDV and SARS-CoV2 (**Fig. 3C & 3D**)^10,45,46^ and used for Co-IP. The results show that N-ΔCTD retained its ability in binding to PKA Cα, while N-ΔNTD did not (**Fig. 3E-3G**). Similarly, only wildtype N protein and N-ΔCTD, but not N-ΔNTD, could enhance the nuclear import of PKA Cα **(Fig. S2)**. These findings clearly demonstrate that the NTD region of PEDV N is responsible for its interaction with PKA Cα and subsequent nuclear localization. To investigate if the N proteins of other coronaviruses are also involved in interacting with PKA Cα, recombinant vectors expressing HA-tagged N proteins of SARS-CoV, SARS-CoV2, MERS-CoV, TGEV and PDCoV were prepared and transfected into the HEK-293 cell. Co-IP clearly showed the interaction of PKA Cα with all the CoV N proteins (**Fig. 3H**).

**Fig. 3.**
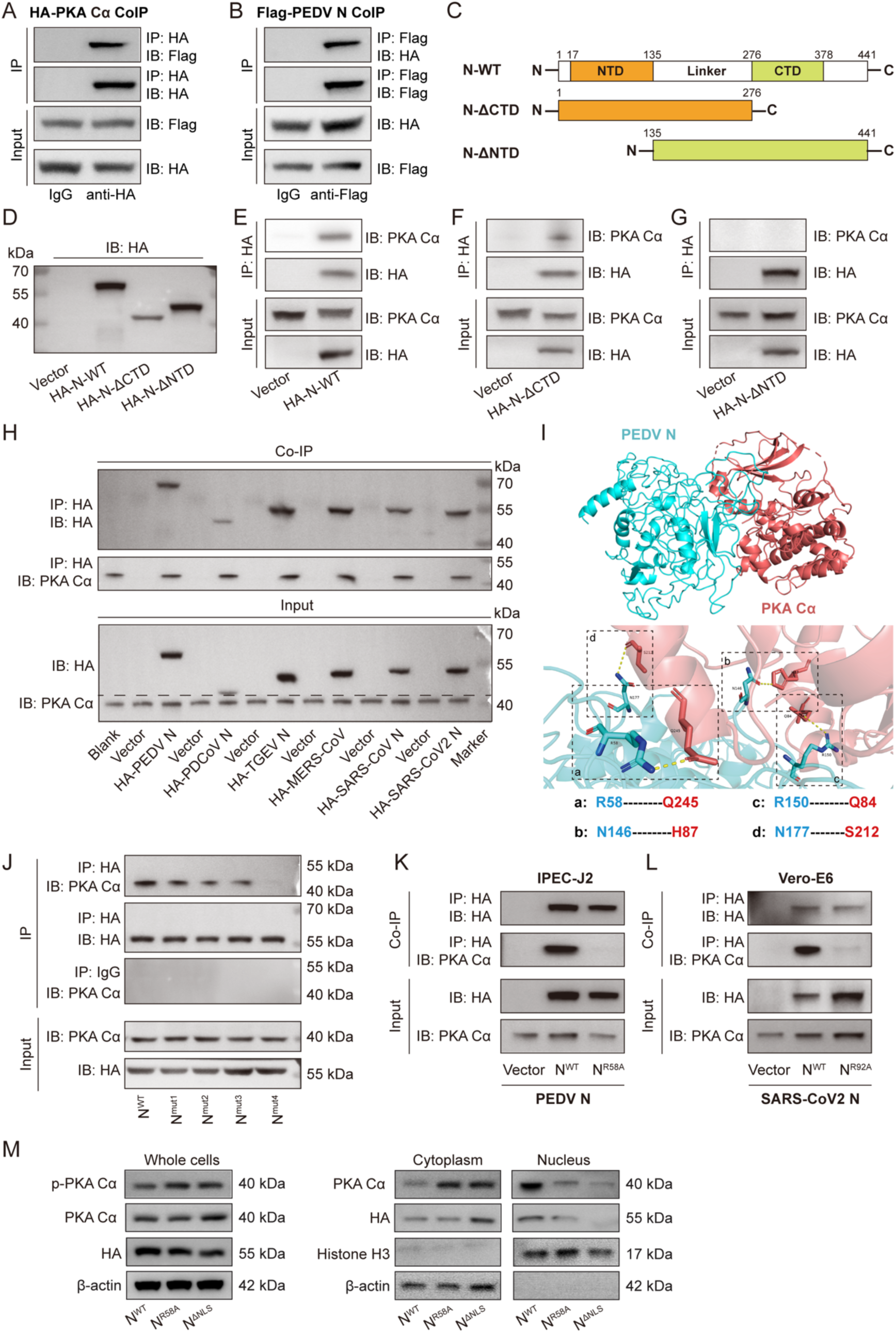
Nucleocapsid proteins of coronaviruses bind to PKA Cα via the conserved arginine within the N-terminal domain. (A)&(B) PEDV N interacts with PKA Cα shown by Western blotting of the immunoprecipitated (IP-WB) samples using anti-HA (A) or anti-Flag antibodies (B) of the lysates of IPEC-J2 cells co-transfected with pCMV-HA-PKA Cα and pCMV-Flag-PEDV N for 24 h. (C) Diagrammatic layout of wildtype PEDV N protein, and (D) expression of the truncated mutants N-ΔNTD and N-ΔCTD in IPEC-J2 cells. (E), (F)&(G) Detection of PKA Cα binding to N (E) and N-ΔCTD (F) but not to N-ΔNTD (G) revealed by IP-WB of the cell lysates. (H) PKA Cα binding to N proteins of different coronaviruses by IP-WB. (I) Structure predication and protein-protein interaction at the interface of PEDV N and PKA Cα. (J) PKA Cα binding to PEDV wildtype N^WT^ and its mutants N^mut1^ (S17A), N^mut2^ (V39A/P40A) and N^mut3^ (G51A/Y52A), but not to N^mut4^ (R58A) shown by IP-WB of the lysates of N-expressing IPEC-J2 cells. (K)&(L) PKA Cα binding to PEDV or SARS-CoV2 wildtype N, but not to their mutant N^R58A^ or N^R92A^. (M) PKA Cα distribution in cytoplasmic and nuclear fractions was distinct between the IPEC-J2 cells expressing PEDV N^R58A^ or N^ΔNLS^ and those expressing wildtype N.

To dissect the molecular basis of the interaction between PKA Cα and N proteins, the crystal structures of PKA Cα and PEDV N as a model were first obtained or simulated by the online website RCSB PDB (https://www.rcsb.org) and I-Tasser (https://seq2fun.dcmb.med.umich.edu//I-TASSER/), respectively. The interface of these two proteins was predicted by H-DOCK (**Fig. 3I** upper, blue for PEDV N, and red for PKA Cα). The prediction obtained four specific interface residue pairs for PEDV N and PKA Cα interaction: R58-Q245, N146-H87, R150-Q84 and N177-S212 (**Fig. 3I** lower). Multiple sequence alignment showed that R58 of PEDV N is highly conserved in different CoV N proteins (corresponding to R54 in PDCoV, R75 in TGEV, R83 in MERS-CoV, R92 in SARS-CoV2, and R93 in SARS-CoV) (**Fig. S3A**), indicating that this residue might play a role in the interaction between different CoV N proteins and PKA Cα. There are also four conserved residues of charged or polar amino acids of different CoV N proteins between S17 and R58 using the PEDV N as the reference sequence (**Fig. S3B upper**) that might also be involved in interacting with PKA Cα. We constructed these four site-specific mutants N^mut1^ (S17A), N^mut2^ (V39A/P40A), N^mut3^ (G51A/Y52A) and N^mut4^ (R58A) on the PEDV N backbone **(Fig. S3B lower)**. **Figure 3J** shows that the N mutants 1, 2 and 3 could still bind to PKA Cα while N^mut4^ did not, suggesting that R58 in PEDV N is the key residues in binding to PKA Cα (**Fig. 3J & 3K**). Mutations of corresponding arginine residues of the N proteins of other CoVs, including SARS-CoV2, also abolished their interaction with PKA Cα (**Fig. 3L & Fig. S3C**). These results demonstrate that this arginine residue in the NTD of different CoV N proteins play a key role in binding to PKA Cα.

Our next questions were if the R58 residue of the N protein essential for the interaction between N protein and PKA Cα is also involved in facilitating PKA Cα entry into the nucleus. We found that the N^R58A^ mutant led to decreased abundance of PKA Cα in the nuclear compartment similar to the N^ΔNLS^ as opposed to the wildtype N protein (**Fig. 3M**), indicating that the N protein depends on the residue R58 for its binding to and nuclear translocation of the PKA Cα via co-mobilization.

### 4. N protein-mediated nuclear entry of PKA Ca dampens its anti-PEDV effect in the cytoplasm

The above results demonstrate that PEDV infection activated the cAMP-PKA signaling and hijacked PKA Cα into the nucleus via its N protein-mediated interaction. Then we wondered if there would be biological significance of the N protein-PKA Cα interaction on the virus. First, we looked at the effects of PKA Cα on PEDV replication by overexpression, knockdown or chemical inhibition in IPEC-J2 cells. Western blotting and/or quantification of viral genome copies showed that PKA Cα overexpression significantly enhanced PEDV replication (**Fig. S4A-S4C)**. By contrast, PKA Cα knockdown attenuated PEDV replication (**Fig. S4D-S4F**). Collectively, these results suggest that PKA Cα appeared to facilitate PEDV replication in IPEC-J2 cells.

As reported previously, PKA Cα phosphorylation at T197 is critical for its activation^47,48^. By overexpression, we found that T197A mutation reduced its nuclear distribution as compared to its wildtype counterpart (**Fig. 4A-4C**). However, nuclear localization of T197A mutant was almost similar to the wildtype PKA Cα in the presence of N protein (**Fig. 4D/d**). Also, T197A mutation enhanced PEDV replication as the wildtype PKA Cα did, as shown by increased N protein expression (**Fig. 4E**). These findings, together with N protein-mediated PKA Cα nuclear translocation (**Fig. 3N**), suggest that PEDV utilizes its N protein to enhance intranuclear localization of PKA Cα independent of T197 phosphorylation to benefit viral replication.

**Fig. 4.**
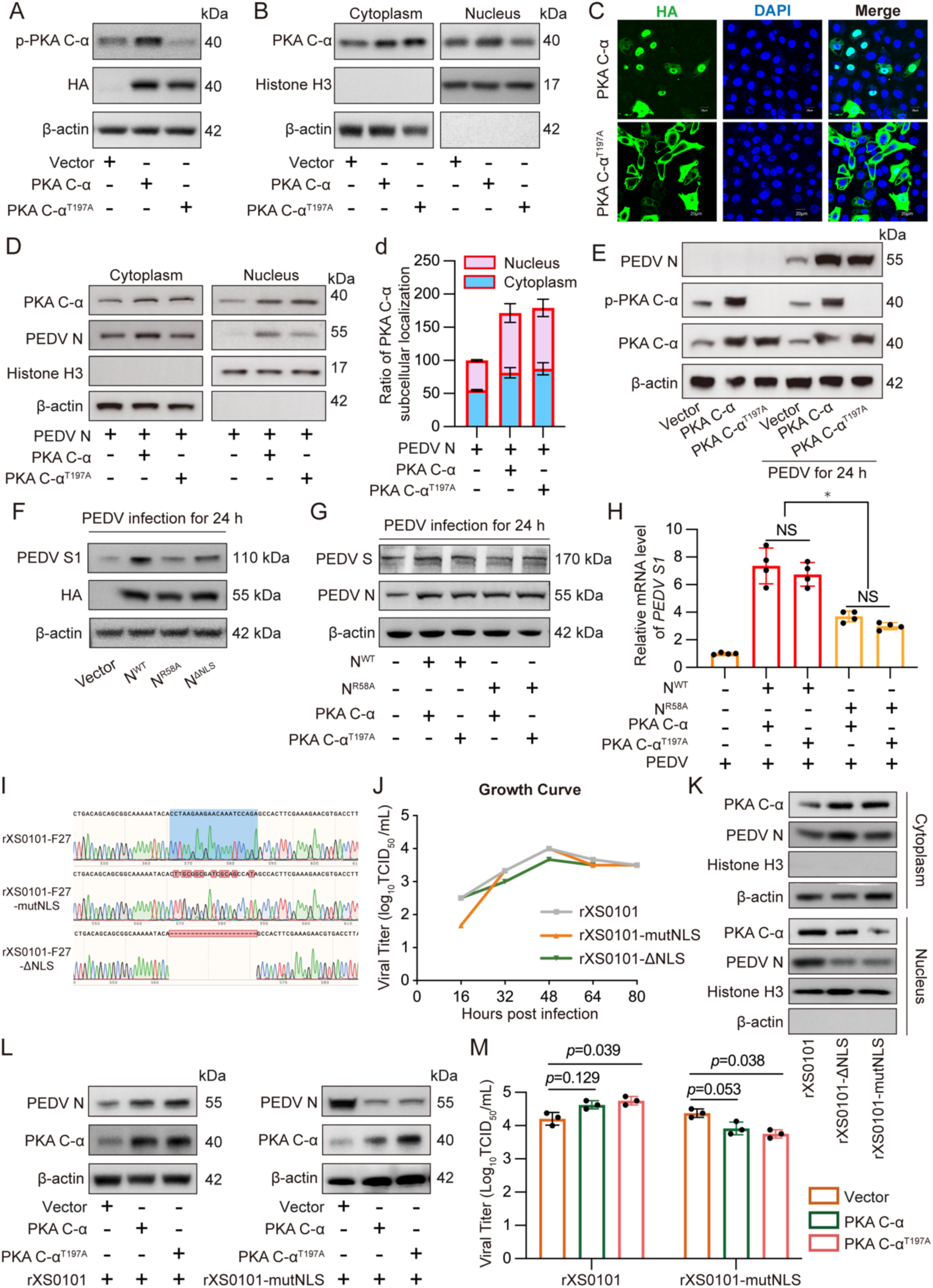
Nucleocapsid protein-mediated nuclear entry of PKA Cα favors PEDV replication. (A)&(B) Phosphorylation and subcellular localization of PKA Cα detected in IPEC-J2 cells expressing PKA Cα and PKA Cα^T197A^. (C) Subcellular localization of exogenous PKA Cα detected by immunocytochemistry using anti-HA antibody. (D)&(d) Distribution of PKA Cα in the nuclear and cytoplasmic fractions of IPEC-J2 cells co-transfected with pCMV-N and pCMV-PKA Cα or -PKA Cα^T197A^. (E) Over-expression of PKA Cα and PKA Cα^T197A^ enhanced PEDV replication shown as increased N in IPEC-J2 cells infected with PEDV for 24 h. (F) Expression of N^R58A^ or N^ΔNLS^ repressed PEDV replication, shown as reduced viral S1 than N^WT^ in the IPEC-J2 cells transfected with pCMV-HA-N^WT^, -N^R58A^ or -HA-N^ΔNLS^ (for 24 h) followed by PEDV infection for another 24 h. (G)&(H) Co-expression of N^R58A^ and PKA Cα or PKA CαT197A inhibited PEDV replication shown by reduced viral S1 protein (G) and S1 mRNA (H) in the IPEC-J2 cells co-transfected with pCMV-HA-N^WT^ or -N^R58A^ and PKA Cα or PKA Cα^T197A^ for 24 h followed by PEDV infection for another 24 h. (I) Sequence verification for mutant PEDV strain rXS0101-mutNLS and rXS0101-ΔNLS. (J) Growth curves of recombinant PEDV strains. (K) Nuclear distribution of PKA Cα was more abundant in IPEC-J2 cells infected for 24 h with the wildtype PEDV than the mutant viruses with NLS mutation or deletion. (L)&(M) Expression of PKA Cα or PKA Cα^T197A^ promoted replication of wildtype PEDV strain rXS0101, but repressed replication of the mutant virus rXS0101-mutNLS as shown by N level (L) and viral titers (M) in IPEC-J2 cells infected for 24 h. The experiments of panels D and H were repeated 3 times for density analysis (d and h). The results of panels h and m were shown as means ± SD, *, *p*<0.05.

To investigate if the N protein mutants (N^R58A^ or N^ΔNLS^) that did not promote PKA Cα nuclear translocation would affect PEDV infection, overexpression of the wildtype N or its mutants in IPEC-J2 cells was approached prior to PEDV infection. The results indicate that PEDV replication was enhanced by wildtype N but diminished by the mutant N^R58A^ or N^ΔNLS^, as shown by PEDV S1 expression and viral genome copies (**Fig. 4F & S5**). Furthermore, we performed the co-expression of N protein/N^R58A^ and PKA Cα/PKA Cα^T197A^ to see their combinatorial effects on PEDV replication. We demonstrated that both wild-type PKA Cα and the PKA Cα^T197A^ mutant enhanced PEDV replication in the presence of wildtype PEDV N protein. However, this pro-viral effect was abolished when the R58 residue was absent (**Fig. 4G & 4H**). These results suggest that the N protein-mediated enrichment of PKA Cα in the nuclear compartment favors viral replication.

To determine if loss of NLS of the N protein would affect PKA Cα partitioning and viral replication at the virus level, we constructed recombinant PEDV viruses containing alanine substitution of the NLS residues (rXS0101-mutNLS) or NLS deletion (rXS0101-ΔNLS) (**Fig. 4I**) that should abolish the N protein-mediated PKA Cα nuclear translocation, hence changing its distribution pattern in the cytosol and nuclei. The NLS mutant viruses showed similar growth to the wildtype recombinant XS0101 strain (rXS0101) (**Fig. 4J**). The NLS mutant viruses led to decreased distribution of both PKA Cα and N protein in the nuclear fraction together with their enrichment in the cytoplasmic compartment in comparison with the wildtype virus strain (**Fig. 4K**). Because nuclear distribution of the PKA Cα, either T197A mutant or wildtype, depends on the localization of the PEDV N protein (**Fig. 4D**), we examined if NLS affects PEDV replication by using the NLS mutant virus in IPEC-J2 cells overexpressing PKA Cα. We found that overexpression of both wildtype PKA Cα and its T197A mutant enhanced virus replication in the IPEC-J2 cells infected with the wildtype PEDV virus rXS0101, but repressed virus replication in the cells infected with the NLS mutant virus (rXS0101-mutNLS) (**Fig. 4L & 4M**). The above results clearly reveal that PEDV utilizes its N protein to escort PKA Cα into the nuclear compartment to favor its replication. However, cytoplasmic enrichment of PKA Cα in the cells infected with rXS0101-mutNLS with reduced viral replication suggest that PKA Cα exerts antiviral effects in the cytoplasm.

### 5. Cytoplasmic PKA Cα induces JAK-independent STAT1 activation and antiviral response which is inhibited by viral N protein-mediated PKA Cα nuclear translocation

Previous studies indicate that CoV N proteins, including those of PEDV and SASR-CoV2, act to evade from innate immune responses by interfering with IFN signaling^49–51^. Type III IFNs are the main players in the innate mucosal immunity of the gut^52–54^. We hypothesized that PKA Cα might exerts its antiviral effects in the cytoplasm by enhancing type III IFN signaling. To investigate whether the cytoplasmic compartment of PKA Cα is capable of directly activating STAT1, overexpression of PKA Cα followed by JAK1/2 inhibitor ruxolitinib or/and H89 (PKA inhibitor) treatment was conducted. As shown in **Fig. 5A**, PKA Cα was involved in rIFN-λ3-induced STAT1 phosphorylation which was counteracted not only by ruxolitinib, but also by H89 treatment, suggesting that PKA might directly activate STAT1 signaling. To further investigate the mechanism by which cytoplasmic PKA Cα mediates STAT1 activation independently of upstream JAK1 activation, we generated four site-directed PKA Cα mutants (K73A^55^, R195A^56^, Y205A^57^, and S339A^58^ in *Sus scrofa*) to identify the critical amino acid residues involved in regulating STAT1 phosphorylation based on previous structural analyses of kinase active sites. **Fig. 5B** shows that only the S339A mutant abolished STAT1 phosphorylation and ISG15 expression compared to wild-type PKA Cα, suggesting that PKA Cα S339 is the critical residue for STAT1 phosphorylation. By combined mutation of T197A and S339A, we confirmed that S339 is the key residue of PKA Cα in activating STAT1 (**Fig. S6)**.

**Fig. 5.**
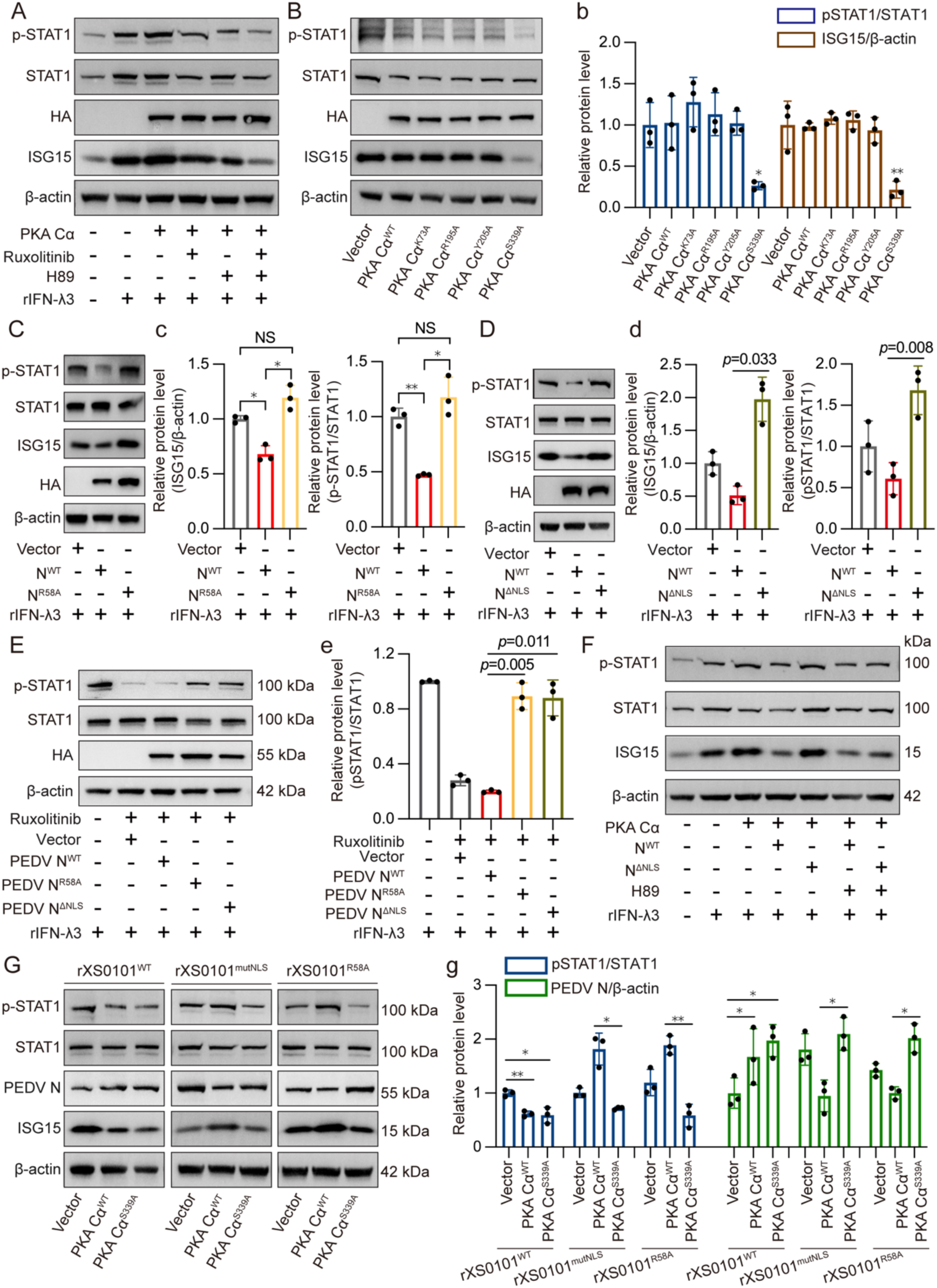
Cytoplasmic PKA induces STAT1 signaling and antiviral response independent of JAK1/2 that is repressible by N protein-mediated promotion of PKA Cα entry into nuclei. (A) Over-expression of PKA Cα enhanced STAT1 phosphorylation inhibitable by PAK inhibitor H98 in the IPEC-J2 cells transfected with pCMV-HA-PKA Cα for 24 h. JAK1/2 specific inhibitor, ruxolitinib, PKA specific inhibitor, H89 and rIFN-λ3 were added at 4 h post transfection. Total STAT1, p-STAT1 and ISG15 expression levels were analyzed by Western blotting. (B) The mutant PKA Cα^S339A^ failed to phosphorylate STAT1 with reduced ISG15 expression in the IPEC-J2 cells expressing HA-tagged PKA Cα or its mutants for 24 h. (b) Densitometric quantification of (B). (C)&(D) The wildtype N inhibited p-STAT1 and ISG15 expression, while the mutant N^R58A^ or N^ΔNLS^ did not. IPEC-J2 cells were transfected with pCMV-HA-N^WT^, -N^R58A^ or -N^ΔNLS^ for 24 h, followed by rIFN-λ3 treatment for next 18 h. (c)&(d) Densitometric quantification of (C)&(D). (E)&(e) p-STAT1 level was less affected in IPEC-J2 cells expressing N^R58A^ or N^ΔNLS^ than the wildtype N by JAK1/2 inhibition. The cells were pretreated with 20 nM ruxolitinib for 30 min, and then transfected with pCMV-HA-N^WT^, -N^R58A^ or -N^ΔNLS^, followed by rIFN-λ3 treatment. (F) p-STAT1 level in IPEC-J2 cells co-expressing PKA Cα and N^WT^ or its mutant N^ΔNLS^ were downregulated by PKA inhibition. The cells were co-transfected with pCMV-HA-PKA Cα and pCMV-N^WT^ or -N^ΔNLS^, followed by H89 and rIFN- λ3 treatments. (G)&(g) Infection of the PKA Cα-expressing IPEC-J2 cells with mutant PEDV rXS0101^mutNLS^ or rXS0101^R58A^, but not the wildtype virus, upregulated STAT1 phosphorylation with reduced viral replication. IPEC-J2 cells were transfected with pCMV-PKA Cα^WT^ or -PKA Cα^S339A^ for 24 h, followed by viruses (rXS0101^WT^, rXS0101^mutNLS^, and rXS0101^R58A^) infection for additional 24 h. The experiments were repeated 3 times for density analysis (g). The results of panels b, c, d, e, and g were shown as means ± SD, *, *p*<0.05; **, *p*<0.01.

Then we used the mutant N proteins with R58A substitution or NLS deletion that would enrich the cytosolic fraction of PKA Cα to see if PEDV employs its N protein to mediate PKA Cα nuclear translocation as a mechanism to evade from STAT1 signaling. We found that IFN-λ3-induced upregulation of pSTAT1 and ISG15 was suppressed by the wildtype N protein but not by N^R58A^ or N^ΔNLS^ mutant protein (**Fig. 5C/c**, **Fig. 5D/d**). Transcription of both *isg15* and *oas1* genes was significantly reduced in the IPEC-J2 cells expressing wildtype N protein, but not by the mutant N^R58A^ (**Fig. S7**). These results indicate that PEDV does utilize its N protein to dampen STAT1 signaling.

Ruxolitinib is a specific JAK1/2 inhibitor which can efficiently suppress downstream STAT1 phosphorylation and ISG15 expression in IPEC-J2 cells (**Fig. S8**). In order to address if PKA Cα mediates STAT1 activation independent of the JAK/TYK2 phosphorylation, we used ruxolitinib in the cells overexpressing wildtype N or N^ΔNLS^/N^R58A^ mutants, followed by IFN-λ3 stimulation. We found that in the cells expressing N^R58A^ and N^ΔNLS^, STAT1 phosphorylation was still enhanced irrespective of JAK inhibition (**Fig. 5E/e**). In order to test our hypothesis that PKA Cα might be hijacked into the nucleus by N protein to interfere with STAT1 activation in the cytoplasm, co-expression of PKA Cα and N^WT^/N^ΔNLS^ were performed. **Figure 5F** shows that PKA Cα could significantly enhance STAT1 phosphorylation and ISG15 expression which could be inhibited by PEDV wildtype N protein but not by N^ΔNLS^, and that H89 treatment inhibited STAT1 activation in the cells co-expressing PKA Cα and N^ΔNLS^. These findings demonstrate that co-mobilization of N protein and PKA Cα into the nuclear compartment is critical for dampening PKA Cα-mediated STAT1 activation.

We then examined if PEDV with NLS mutation and R58 substitution in the N protein would affect STAT1 activation and PEDV replication in IPEC-J2 cells and we attempted to dissect if N protein nuclear shuttling or its binding with PKA Cα is involved in PKA Cα-mediated STAT1 activation in the PEDV-infected cells. **Figure 5G/g** shows that STAT1 activation was downregulated in the wildtype PEDV-infected cells expressing either wildtype PKA Cα or mutant PKA Cα^S339A^. However, in the cells infected with the mutant virus rXS0101^mutNLS^ or rXS0101^R58A^, only the mutant PKA Cα^S339A^ expression inhibited STAT1 activation while the wildtype PKA Cα did not. Downregulation of STAT1 activation is accompanied with increased viral replication in all these instances (**Fig. 5G/g**). These results illustrate that PKA Cα is involved in STAT1 activation in the cytoplasm of virus-infected cells, nuclear mobilization of PKA Cα is essential to diminish STAT1 activation, and PEDV deploys its N protein to evade from PKA Cα-mediated antiviral innate immunity by hijacking PKA Cα into the nuclear compartment.

## Discussion

The mammalian hosts can sense the invading viruses via the pattern recognition receptors (PRR), such as Toll-like receptors (TLRs), cyclic GMP-AMP synthase (cGAS), etc., to initiate the antiviral innate immune responses via a host of sophisticated signaling pathways^59^, while human and animal coronaviruses have developed multiple strategies to evade host innate immunity^60,61^. Majority of the coding proteins in coronaviruses have been reported to antagonize the host innate immune system by either targeting viral sensors or blocking downstream antiviral signaling molecules^60,61^. However, it remains unknown if the cAMP-protein kinase A signaling is involved in activation of the innate immune responses during viral infections, including those of coronaviruses, and whether this pathway is manipulated by the viruses or viral proteins. Here we report for the first time that the N proteins of coronaviruses activated the cAMP-ADCY10-PKA signaling in IPEC-J2 or VERO-E6 cells, cytoplasmic PKA activates STAT1 and ISG15 expression independently of JAK, and coronavirus N proteins promotes PKA Cα into the nuclei by direct interaction via a specific arginine residue conserved in coronaviruses to escape from PKA-mediated STAT1 activation (**Fig. 6**).

**Fig. 6.**
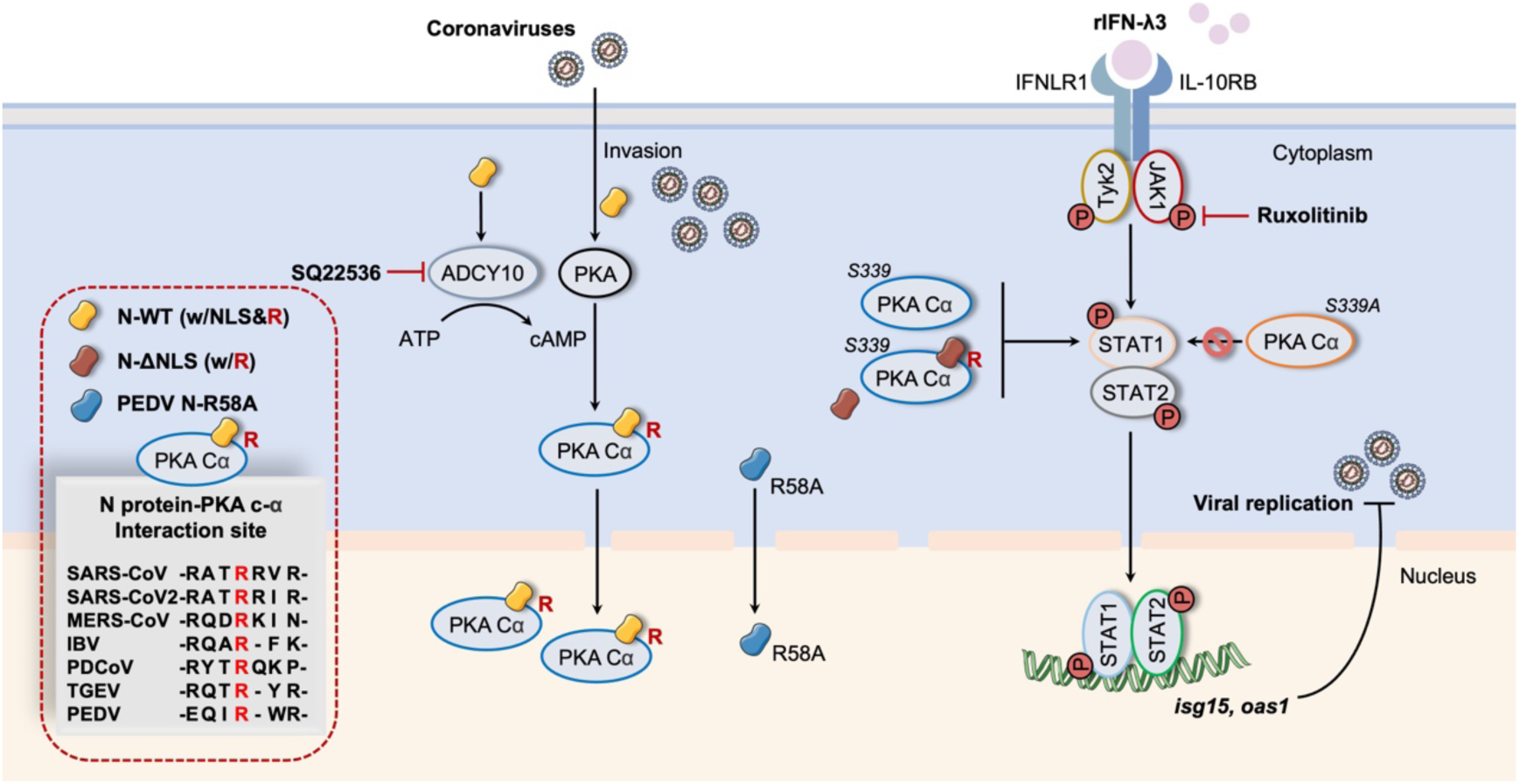
Schematic model depicting the role of PKA in phosphorylating STAT1 in cytoplasm and the part that nucleocapsid proteins of coronaviruses play in antagonizing PKA-STAT1 signaling. Infection of PEDV, as a model coronavirus, upregulates expression of soluble ADCY10 via its N protein with increased cAMP production and PKA activation, which can be counteracted by specific ADCY inhibitor SQ22536. Cytoplasmic PKA Cα, with Ser339 as its key residue for the kinase activity, upregulates STAT1 phosphorylation independently of IFN-λ3-induced JAK/STAT1 signaling. However, N proteins of the coronaviruses interact with PKA Cα via the conserved arginine, R58 of PEDV N, R92 of SARS-CoV2 N, etc., and hijack PKA Cα into the nucleus via their NLS-mediated nuclear translocation. Thus, coronavirus N proteins intercept PKA Cα-mediated STAT1 activation in the cytoplasm and suppress expression of downstream ISGs in favor of viral replication.

The cAMP-PKA pathway plays a key role in regulation of a plethora of cellular functions in almost all tissues in mammals, such as cell proliferation, gene expression, apoptosis and several metabolic processes^30,62^. The regulatory (R) subunits control PKA activity depending on the levels of the second messenger cAMP. In the absence of cAMP, the R subunits bind to the catalytic (C) subunits and suppress their activity, while binding of two molecules of cAMP to each R subunit leads to conformational change in the holoenzyme, releasing the C subunits to phosphorylate downstream targets^35,63^. So far, there has been no study demonstrating that STAT1 is the substrate of PKA for its activity. Initially, we found that STAT1 phosphorylation was repressible by PKA inhibitor H89 in a level similar to the JAK inhibitor ruxolitinib in the rIFN-λ3-treated IPEC-J2 cells expressing PKA CαT197A (**Fig. 5A**), suggesting that STAT1 might be the substrate of PKA. Four additional site-directed PKA Cα mutants K73A^55^, R195A^56^, Y205A^57^, and S339A^58^ in *Sus scrofa* known to be involved in PKA activity as well as a double mutant PKA Cα^T197A/S339A^ were generated to identify the critical amino acid residues involved in regulating STAT1 phosphorylation. We demonstrated that downregulation of STAT1 phosphorylation was more significant with the S339A mutant PKA Cα than the other mutants, including the T197A mutant (**Fig. 5B & 5C**). Although the conserved activation loop site T197 phosphorylation is known to stabilize and activates PKA^64^, Keshwani et al reported that S338 (S339 in *Sus scrofa*) phosphorylation is required for solubilization of the S subunit and subsequent T197 phosphorylation^58^. Therefore, we conclude that PKA is involved in positive regulation of STAT1 activation independent of the JAK.

The N proteins of the coronaviruses, critical for viral genome encapsulation and replication by binding to viral genome RNA^65^, are known to interfere with IFN signaling to evade host antiviral responses. SARS-CoV N protein suppresses IFN-β production by direct interaction with host protein activator of protein kinase R (PACT) to disrupt its binding with RIG-I/MDA5^66^. MERS-CoV N protein could inhibit both RIG-I induced types I and III IFN induction by interacting with host TRIM25 to restrain RIG-I ubiquitination^67^. The N protein of porcine delta coronavirus (PDCoV), another emerging swine coronavirus, antagonizes IFN-β production by interacting with RIG-I and MDA5^68^. The PEDV N protein inhibits IFN-β production and ISG expression by targeting the TBK1 to interfere its interaction with IRF3^69^. Recent work in our laboratory reveals that PEDV N could inhibit host antiviral responses by binding to the transcriptional factor Sp1 to downregulate HDAC1 transcription, leading to STAT1 acetylation over phosphorylation^24,25^. In the current study, we have identified that PKA-Cα is a novel target of the N proteins of coronaviruses to escape the host IFN-induced responses. We elaborate that the specific arginine residue at 58 of PEDV N (and R54 of PDCoV N, R75 of TGEV N, R83 of MERS CoV, R93 of SARS-CoV, and R92 of SARS-CoV2) is responsible for the interaction of coronavirus N proteins with PKA Cα (**Fig. 3**; **Fig. S3**) and such interaction is critical to reduce the PKA Cα level in the cytoplasmic compartment by promoting its nuclear translocation, hence the downregulation of its activity of STAT1 phosphorylation (**Fig. S9**). This finding is supported by the fact that NLS deletion mutant N^ΔNLS^ also increased cytoplasmic PKA Cα (**Fig. 3M**) and upregulated STAT1 phosphorylation (**Fig. 5E**) similar to the mutant N^R58A^. This novel mechanism is distinct from the study by Mu et al (2020) who reported that SARS-CoV-2 N protein antagonizes IFN-I signaling by interaction with STAT1 and STAT2, hence inhibition of their activation^19^.

While there are significant differences in the amino acid sequences of the N proteins of different coronaviruses, there are several major conserved sites within the N-terminal domain (NTD) of their N proteins. For instance, the conserved R95 residue in the N protein of SARS-CoV2 is methylated by host arginine methyltransferase 1, and R95 methylation inhibits stress granules formation and facilitates viral replication^70^. We have demonstrated that the R58 residue of the PEDV N protein is a key site for interaction with PKA Cα. Sequence alignment reveals that this R58 residue is conserved in the NTDs of other coronaviruses: R92 of SARS-CoV2, R93 of SARS-CoV, R83 of MERS-CoV, R75 of TGEV, and R54 of PDCoV that are also involved in interaction with PKA Cα (**Fig. S3**) and downregulation of STAT1 signaling (**Fig. S9**). To investigate the role of R58A in STAT1 activation and viral replication, PEDV was used as a model coronavirus to construct mutant viruses. We found that R58A mutant virus rXS0101^R58A^ did exhibit reduced replication similar to rXS0101-mutNLS, as compared to the wildtype virus. Upregulation of PKA Cα-mediated STAT1 activation and ISG15 expression might account for suppression of these mutant viruses in IPEC-J2 cells because expression of the mutant PKA Cα^S339A^ showed reduced kinase activity on STAT1 phosphorylation and increased viral replication (**Fig. 5G**).

All the above results demonstrate that PEDV employs its N protein to promote nuclear translocation of PKA Cα via direct R58-mediated interaction to reduce the abundance of PKA Cα in the cytosol where it phosphorylates STAT1 to enhance the antiviral immune response. This is apparently alternative to the canonical JAK-STAT1 signaling. We propose that this mechanism also exist in other coronaviruses, such as SARS-CoV-2, MERS-CoV, etc. Reverse genetic approach could be used in BL3 laboratories to confirm if PKA-STAT1 signaling and its interference by the N proteins also operates in these emerging human coronaviruses.

Our study provides evidence that PEDV infection regulates ADCY10-cAMP signaling. PEDV and its N protein significantly upregulated expression of *ADCY10* encoding the soluble adenylate cyclase 10 (sADCY10), leading to increased cytoplasmic cAMP and PKA Cα activation (**Fig. 1**). The N proteins of other coronaviruses SARS-CoV2, SARS-CoV, MERS-CoV, TGEV and PDCoV also increased *ADCY10* expression and cAMP production (**Fig. 1L & S1**). Thus, we could postulate that the N proteins of coronaviruses have double targets in the cAMP-PKA signaling: N/ADCY10 and N/PKA. So far there have been no studies on virus-mediated transcriptional activation of ADCYs, including sADCY10. The underlying mechanism of transcriptional activation of sADCY10 expression by N proteins of coronaviruses awaits further investigation.

PKA is found to promote infections of several viruses *in vitro*. For example, PKA plays a role in herpes simplex virus growth by phosphorylating the nuclear ICP4 protein, and viral growth was significantly reduced in the PKA-deficient cells^71^. The endogenous PKA activity is necessary for Zika virus (ZIKV) replication, while the PKA-specific inhibitor, PKI 14-22, inhibits viral infection shown as suppression of negative-sense RNA synthesis and viral protein translation^72^. PKA is beneficial to HIV-1 replication by blocking poly(I:C) and CpG DNA induced *ifn*-α and *ifn*-β gene transcription^73^. Cartier et al reported that PKA C is incorporated into the HIV-1 virus particle that might regulate viral infectivity by interaction with and phosphorylate the capsid protein^74^. We found that PKA Cα, when over-expressed, is also a positive regulator in PEDV infected IPEC-J2 cells. Because over-expressing PKA Cα promote replication only of the wildtype PEDV where the viral N protein is able to co-mobilize PKA Cα into the nuclei, but not of the mutant PEDV rXS0101-mutNLS where the viral N protein fail to promote PKA Cα’s entry into the nuclei (**Fig 4L & 4M; Fig. S4**), we propose that PKA Cα in the nuclear compartment contributes to increased PEDV replication. Further research is to be pursued to examine the molecular mechanism with which nuclear PKA Cα acts to enhance replication of PEDV or possibly other coronaviruses.

In conclusion, we have elaborated the novel mechanism of PKA Cα-mediated STAT1 activation in the cytoplasmic compartment in PEDV-infected cells which is distinct from the canonical JAK-STAT signaling and could be dampened by the N proteins of coronaviruses via direct interaction with and co-mobilization of PKA Cα into the nuclei (**Fig. 6**). Together with the findings of increased transcriptional expression of *ADCY10* and subsequent cAMP production by coronaviral N proteins as well as enhanced viral replication by nuclear PKA, we anticipate that PEDV could be used as a surrogate coronavirus to decipher the holistic mechanisms and integrated roles of coronaviruses’ N proteins in tilting the balance between the antiviral ADCY10-cAMP-PKA-STAT1 signaling in the cytosol and PKA-mediated proviral effect in the nuclei. Elaboration of such mechanisms could provide multiple novel targets for development of anti-coronaviral drugs.

## Acknowledgments

This work was financially supported by grants from the National Natural Science Foundation of China (Nos. 32372978, 32202771), the “Pioneer” and “Leading Goose” Research and Development Program of Zhejiang Province (No. 2023C02036), the science and technology innovation leading talents project of “High-level Talents Special Support Plan” of Zhejiang Province (No. 2021R52041), and the Analysis and Measurement Foundation of Zhejiang Province (No. LTGC24C080001).

## Author Contribution

J.X., W.F., and X.L. conceived and designed the experiments. J.X. and X.L. wrote the manuscript. J.X. performed the majority of the experiments, data collection and analysis. Y.X., Q.G. and X.X. contributed to virus preparation. W.L. and J.X. contributed to gene knockdown and overexpression. J.X., Y.X., W.L., Z.S., L.Z., Q.G., X.X., Y.S., F.H., L.Z., and X.L. contributed to experimental suggestions and revised the manuscript. All authors approved the final version of the manuscript.

