## Supplemental Figures for "Coronavirus Nucleocapsid Proteins Hijack Host Protein Kinase A Catalytic Subunit α into Nucleus to Evade from STAT1 Signaling"

**Fig. S1. Transcription of the *ADCY* genes in response to expression of N proteins of different coronaviruses in Vero-E6 cells.**

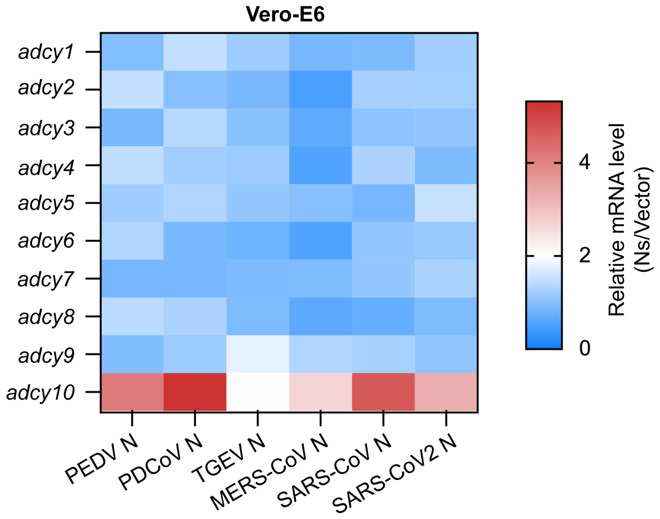

**Fig. S2. The effects of wildtype PEDV N protein and its NTD or CTD deletion mutant on PKA Cα cellular distribution.**

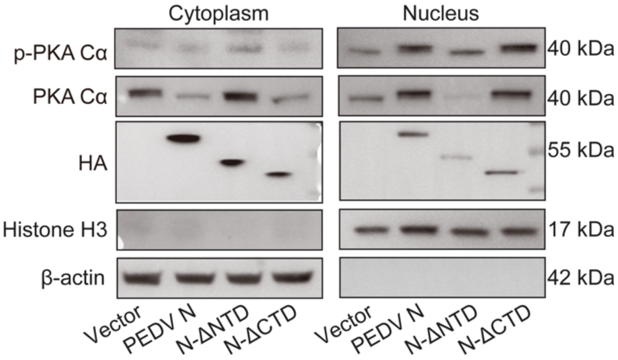

### Fig. S3. The conserved arginine residue of N proteins of several coronaviruses involved in interaction with PKA Cα.

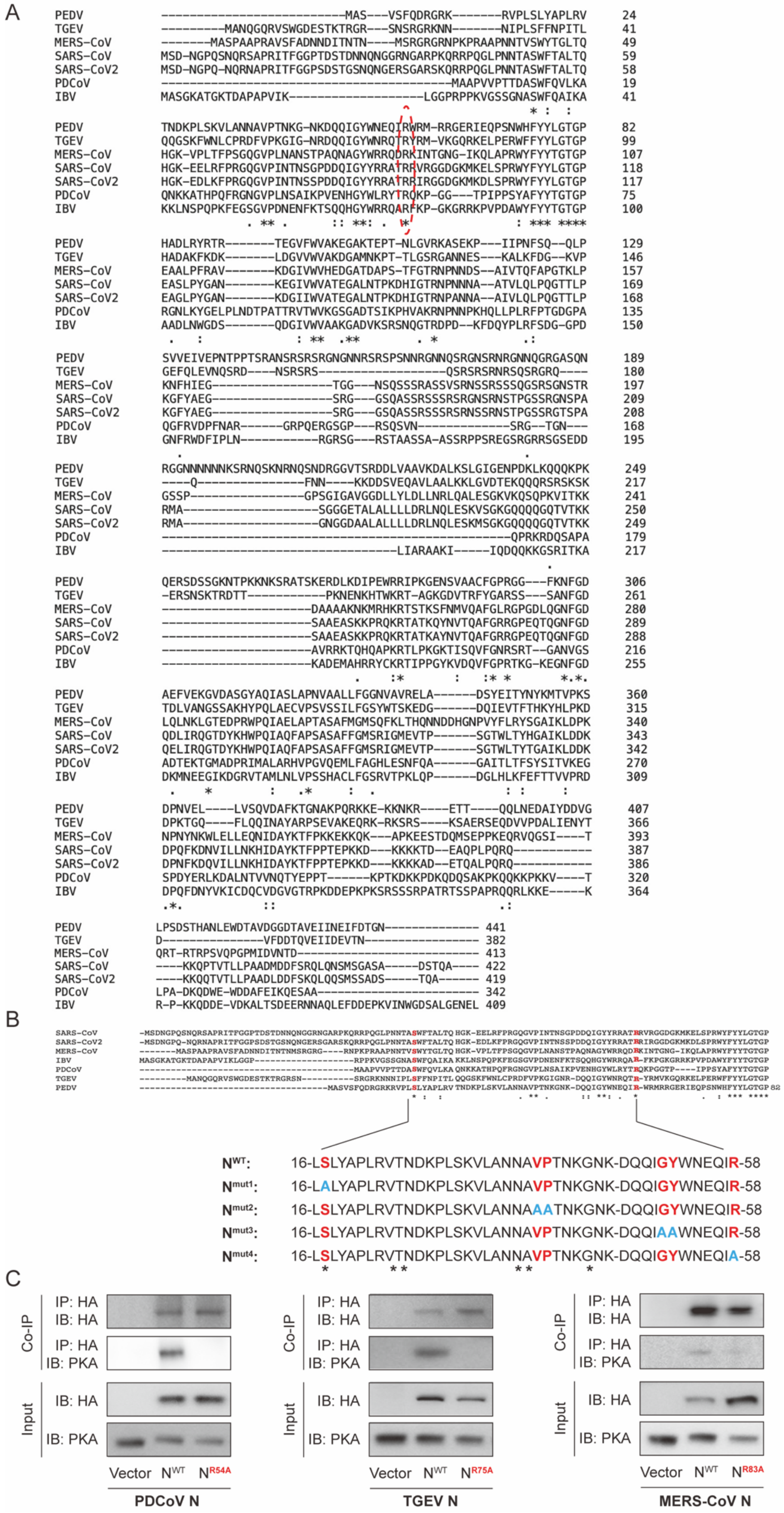

**Fig. S4.** **PKA C-α positively regulates PEDV replication.**

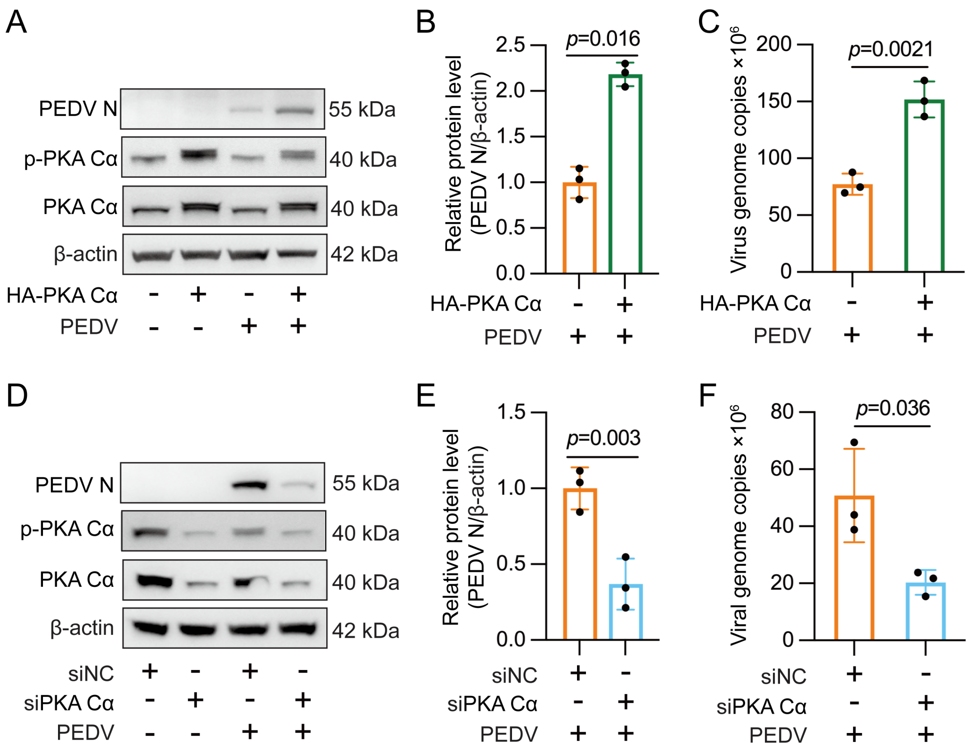

**Fig. S5.** **Both of R58 and NLS of the N protein are critical for PEDV replication.**

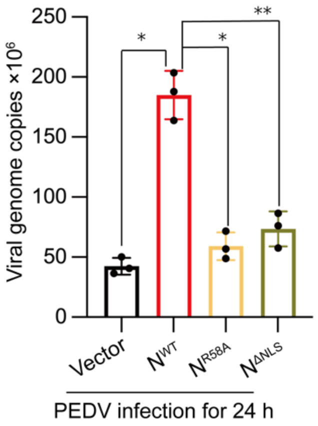

**Fig. S6.** **S339 is the key residue of PKA Cα in activating STAT1.**

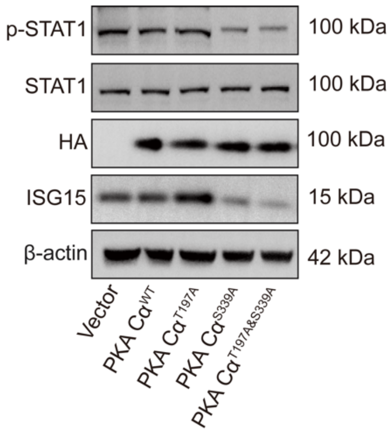

**Fig. S7.** **R58 of PEDV N protein is involved in inhibition of *ISG15* and *OAS1* transcription.**

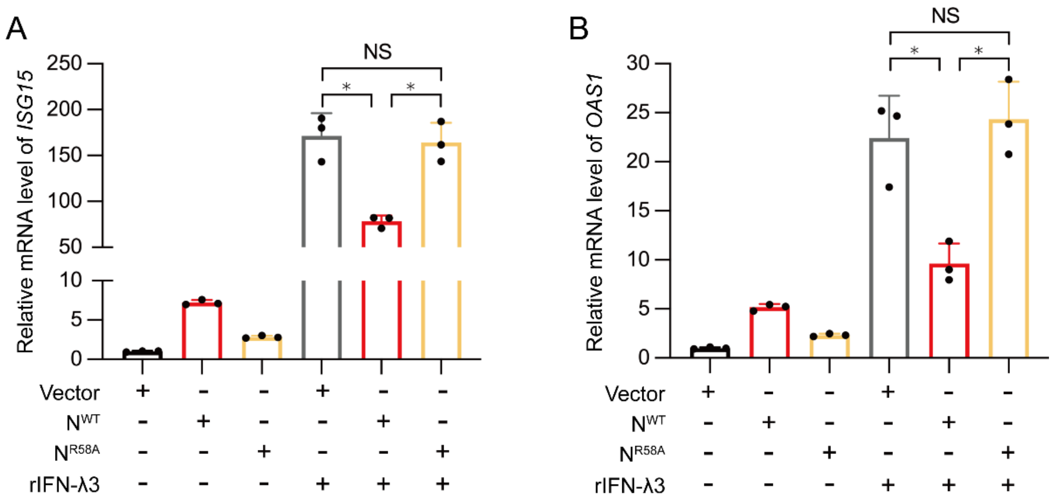

**Fig. S8. The JAK1/2 specific inhibitor Ruxolitinib suppresses downstream STAT1 activation, and PKA Cα does not interact with JAK1.**

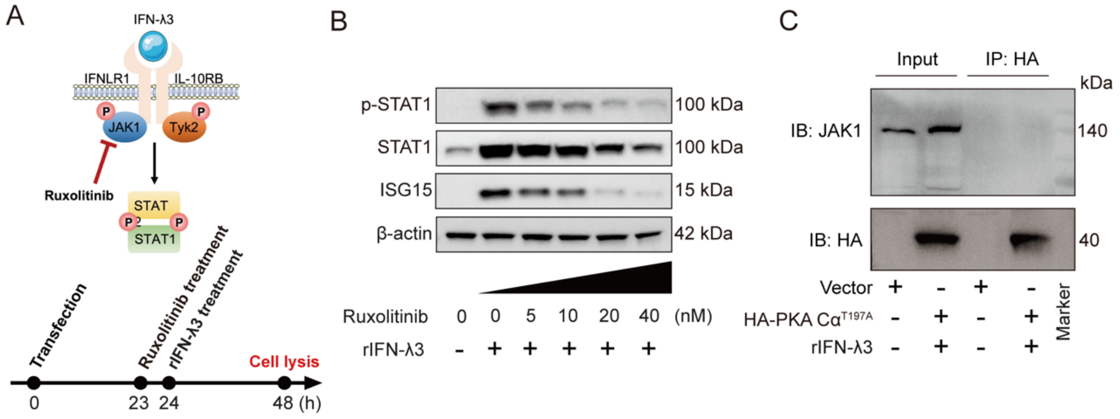

**Fig. S9. The conserved arginine residue in the N proteins of different coronaviruses is involved in STAT1 phosphorylation and ISG15 expression.**

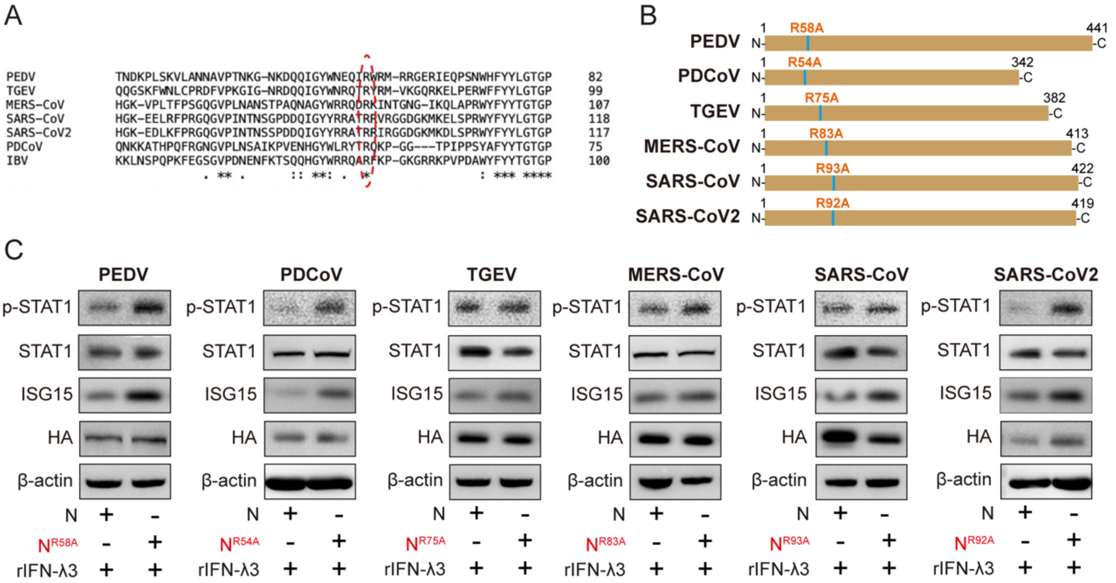
